# Nitrosative stress under microaerobic conditions triggers inositol metabolism in *Pseudomonas extremaustralis*

**DOI:** 10.1101/2023.10.08.561431

**Authors:** Esmeralda C. Solar Venero, Lucia Giambartolomei, Ezequiel Sosa, Darío Fernández do Porto, Nancy I. López, Paula M. Tribelli

## Abstract

Bacteria are exposed to reactive oxygen and nitrogen species that provoke oxidative and nitrosative stress which can lead to macromolecule damage. Coping with stress conditions involves the adjustment of cellular responses, which helps address metabolic challenges. In this study, we performed a global transcriptomic analysis of the response of *Pseudomonas extremaustralis* to nitrosative stress, induced by S-nitrosoglutathione (GSNO), a nitric oxide donor, under microaerobic conditions. The analysis revealed the upregulation of genes associated with inositol catabolism; a compound widely distributed in nature whose metabolism in bacteria has aroused interest. The RNA-seq data also showed heightened expression of genes involved in essential cellular processes like transcription, translation, amino acid transport and biosynthesis, as well as in stress resistance including iron-dependent superoxide dismutase, alkyl hydroperoxide reductase, thioredoxin, and glutathione S-transferase in response to GSNO. Furthermore, GSNO exposure differentially affected the expression of genes encoding nitrosylation target proteins, encompassing metalloproteins and proteins with free cysteine and /or tyrosine residues. Notably, genes associated with iron metabolism, such as pyoverdine synthesis and iron transporter genes, showed activation in the presence of GSNO, likely as response to enhanced protein turnover. Physiological assays demonstrated that *P. extremaustralis* can utilize inositol proficiently under both aerobic and microaerobic conditions, achieving growth comparable to glucose-supplemented cultures. Moreover, supplementing the culture medium with inositol enhances the stress tolerance of *P. extremaustralis* against combined oxidative-nitrosative stress. Concordant with the heightened expression of pyoverdine genes under nitrosative stress, elevated pyoverdine production was observed when *myo*-inositol was added to the culture medium. These findings highlight the influence of nitrosative stress on proteins susceptible to nitrosylation and iron metabolism. Furthermore, the activation of *myo*-inositol catabolism emerges as a protective mechanism against nitrosative stress, shedding light on this pathway in bacterial systems, and holding significance in the adaptation to unfavorable conditions.

## Introduction

Bacterial survival and adaptability in changing environments depend on different resistance mechanisms against several stress agents. Free radicals are highly reactive species, mainly originated from energy producing processes like aerobic and anaerobic respiration, causing damage to different macromolecules. Reactive oxygen species (ROS) are produced during aerobic respiration, while anaerobic respiration, utilizing nitrate as electron acceptor, generates reactive nitrogen species (RNS). ROS and RNS trigger oxidative or nitrosative stress, respectively. RNS include nitric oxide (NO) and its derivatives peroxynitrite (ONOO^−^), nitrosothiols formed through reaction with thiol groups, and nitrotyrosine from nitration of tyrosine by NO, ONOO^−^ or NO_2_^-^ [1–3]. Nitrosative and oxidative stress can also be induced from various environmental compounds, like pollutants, xenobiotics, and antimicrobials, and from UV radiation [4]. NO is also produced by the mammalian immune system and acts as a modulator molecule in both mammals and plants [5–7]. Nitrosative stress can lead to DNA and macromolecular damage, including protein oxidation. Some residues or structures are more susceptible to the RNS or ROS attack such as iron-sulfur (Fe-S) clusters, metal centers, tyrosine and cysteine residues, and thiols [8]. The radical attack to these residues or clusters results in both reversible and irreversible damage, leading to alterations in function and structure[9].

*Pseudomonas* species exhibit a wide variety of energy-generation metabolisms spanning from aerobic to anaerobic, involving NO_3_^-^ reduction, complete denitrification process to N_2_ and even fermentation of arginine or pyruvate [10,11]. These energy-generating pathways enable *Pseudomonas* species to thrive in diverse environmental niches, including soil, water, and host-associated habitats, and they play an integral role in the ecological and physiological success of these microorganisms. *Pseudomonas extremaustralis* is an Antarctic bacterium, capable to growth under different temperatures and oxygen availability. The denitrification process in this bacterium is incomplete, due to the absence of *nir* genes which encode the enzymes required for the reduction of NO_2_ ^-^ to NO, but interestingly, harbors all the *nor* and *nos* genes associated with nitric and nitrous oxide reduction, respectively [12].

Bacterial adaptation to different stress conditions entails changes leading to solve metabolic and structural challenges, such us alterations or damage in DNA, RNA and proteins and envelopes, among others. Some strategies can involve the utilization of alternative pathways to exploit different carbon sources. Thus, when *P. putida* was grown at 10°C, the uptake and assimilation of branched-chain amino acids alongside the activation of the 2-methylcitrate pathway to generate succinate and pyruvate has been identified as mechanism employed to cope with stress when central metabolism is downregulated[13]. Likewise, in cold conditions genes involved in primary metabolism were downregulated in *P. extremaustralis* whereas genes linked to ethanol oxidation were activated, showing that this secondary pathway resulted essential for cold growth [14]. In *P. aeruginosa* has been reported that under oxidative stress the glyoxylate shunt, an alternative to the tricarboxylic acid cycle, increases bacterial survival allowing the utilization of acetate and fatty acids as carbon sources [15].

Considering the importance of RNS derived from both anaerobic respiration and other biotic and abiotic environmental reactions, we evaluated the global transcriptomic response of *P. extremaustralis* to nitrosative stress under low O_2_ tensions using S-nitrosoglutation (GSNO), as a NO donor compound. Our findings show that exposure to GSNO under microaerobic conditions prompted adjustments in central metabolic pathways including iron metabolism along with the upregulation of genes associated with the inositol catabolism. Notably, *myo*-inositol arises as a carbon source supporting *P. extremaustralis* growth and increased nitro-oxidative stress resistance and pyoverdine production. These findings highlight the versatility and adaptability of *P. extremaustralis* to withstand nitrosative stress, by displaying an alternative metabolism, thus contributing to the understanding of the survival mechanisms employed by microorganisms in extreme environments.

## Materials and methods

### Strains and culture conditions

*P. extremaustralis* 14-3b (DSM 25547), a species isolated from the Antarctica [16] was used through the experiments. Bacterial cultures were grown in Lysogeny Broth medium (LB) at 30 °C. Microaerobic cultures were incubated in sealed bottles with a 1:2 medium-to-flask volume ratio and 50 rpm agitation. Microaerobic culture’s medium was supplemented with 0.8 g/l KNO_3_.

GSNO effect on *P. extremaustralis*’ survival and growth was determined in microaerobic cultures exposed to 1, 10 and 100 µM of GSNO. Bacterial growth was evaluated by OD_600nm_ measurement after adding the different concentrations of GSNO at T=0. Optical density was monitored over 24 h at 30min intervals using an automated plate reader (BMG OPTIMA FLUOstar). For survival experiments, cultures were grown for 24h and further exposed to the different concentrations of GSNO for 1h. Afterwards, appropriate dilutions of control cultures (m-C) or cultures exposed to GSNO were plated in LB agar and incubated at 30°C. Colony-forming units per ml (CFU / ml) was determined and the survival percentage was calculated with respect to control cultures.

### RNA extraction and RNA library preparation

Cultures were microaerobically grown in LB medium for 24 h and then incubated for 1 h with 100 µM S-nitrosoglutatione (GSNO, Sigma Aldrich) (m-NS condition) or with the addition of sterile water (m-C). Total RNA was isolated from 6 ml of *P. extremaustralis* m-C and m-NS cultures using the Trizol method. Samples were treated with DNAse I and were validated using an Agilent 2100 Bioanalyzer (Agilent Technologies). To improve the quality of the readings, ribosomal RNA was depleted from the samples, using the RiboZERO Kit (Illumina), following the manufacturer’s instructions. Libraries were prepared using the TruSeq RNA Library Prep Kit v2 (Illumina). Mass sequencing was performed using NextSeq 550 platform with a single-end protocol. For each condition duplicated independent RNA extraction and libraries were used.

### RNA-seq data analysis

Reads were preprocessed using the Trimmomatic tool [17] by eliminating adapters and low-quality sequences. Reads’ quality was evaluated using the Fast QC tool (www.bioinformatics.babraham.ac.uk/projects/fastqc/).

Reads alignment and assembly to the *P. extremaustralis* genome, transcript identification and abundance quantification was carried out using the Rockhopper software that use the Bowtie2, Bayesian and Anders and Huber approaches [18].Reads were normalized per kilobase per million mapped reads (RPKM). Differential gene expression was considered only with PL<L0.05 and QL<L0.05. Concordance between the independent replicates for each of the analyzed conditions was verified by performing a Spearman correlation analysis of normalized counts (Fig. S1).

Genes were assorted into functional classes using KEGG [19], MetaCyc [20] and String [21] tools.

### Prediction of oxidation and nitrosylation targets

The Target-Pathogen tool [22] was used to identify proteins with possible oxidation and / or nitrosylation target sites (Free Cys or Tyr residues or metal biding clusters) within the *P. extremaustralis* genome.

### Comparative genome analysis

General genome sequence analysis was performed using the bioinformatics tools available on National Center for Biotechnology Information (www.ncbi.nlm.nih.gov), including BLAST (Basic Local Alignment Search Tool) [23]. We also used RAST (Rapid Annotation using Subsystem Technology) server [24] and Pseudomonas Genome Database (Pseudomonas.com, [25]). Genomes used, with corresponding GenBank accession numbers, were: *P. extremaustralis:* 14-3b (AHIP00000000.1), USBA 515 (FUYI01), DSM 17835^T^ (LT629689.1), 2E-UNGS (CP091043.1), CSW01(JAQKGS01),1906 (JARIXU01), NQ5 (JARBJR01) strains, *P. protegens* Pf-5 (CP000076.1), and *P. syringae* pv *syringae* B728a (CP000075.1)

### Quantitative Real Time PCR Experiments (RT qPCR)

Total RNA of *P. extremaustralis* microaerobic and m-NS cultures was isolated using the Total RNA Extraction Kit (RBC Biosciences). After treatment with DNaseI, cDNA was obtained using random hexamers (Promega) and Revert Aid reverse transcriptase (ThermoFisher Scientific) according to the manufacturer’s instructions. RT qPCR was performed using a MyiQ2 Real-Time PCR Detection System (Bio-Rad Laboratories, Hercules, USA) and Real-Time PCR mix (EvaGreen qPCR Mix Plus no ROX). The expression of target genes was evaluated using the following primers: *motB* forward 5′ CAACGCCAGCAACAAAGACA 3′ and reverse 5′ CTTCGGAAAACCGCGCATC 3′; *hbo* forward 5′ CGAGCCTAAGAGCAACACCA 3′ and reverse 5′TGAATAGGCTGTCGGCACTG 3′ and *flaB*forward 5’ CGTAACCAGCGCTGACATGGCTC 3’ and reverse 5’ GACAGGATCGCAGTGGAAGCCGA 3’. Expression of the 16S rRNA gene was used for normalization of target genes expression levels in each condition, obtained using primers forward 5′ AGCTTGCTCCTTGATTCAGC 3′ and reverse 5′ AAGGGCCATGATGACTTGAC’ 3. The cycling conditions were as follows: denaturation at 95 °C for 15 min, 40 cycles at 95 °C for 25 s, 60 °C for 15 s, and 72 °C for 15 s, with fluorescence acquisition at 80 °C in single mode. The 2^-ΔΔCT^ method [26] was used to calculate the relative fold gene expression of individual genes.

### Bacterial growth using *myo*-inositol as sole carbon source

*Myo*-inositol metabolism was evaluated in E2 medium [27] supplemented with 10 g/l of *myo*-inositol or glucose under aerobic and microaerobic conditions. Microaerobic cultures were performed as described above while for aerobic experiments, cultures were incubated in Erlenmeyer flasks using 1:10 medium-to-flask volume ratio and 200 rpm agitation. Aerobic and microaerobic cultures were incubated for 24h and 48h, respectively.

Biofilm formation was assayed in E2 medium supplemented with *myo*-inositol or glucose and KNO_3_ in polystyrene microtiter plates with an initial OD_600nm_ of 0.025. After 48h of static incubation at 30°C, OD_600nm_ of planktonic cells (Planktonic cells absorbance, PCA) was measured and biofilms were quantified using the standard crystal violet method [28]. Briefly, attached cells were stained with 200 μl of 0.1% crystal violet, further washed and the colorant was solubilized with absolute ethanol. Crystal violet solution was transferred to flat bottom microtiter plates and OD_570nm_ (Crystal violet absorbance CVA) was measured with a BMG OPTIMA FLUOstar microplate reader. Biofilm formation index was defined as CVA/PCA.

### Pyoverdine production

Pyoverdine production of microaerobic cultures was analyzed in iron-limited E2 medium, without microelements, supplemented with *myo*-inositol or glucose as carbon source and KNO_3._ After 48h pyoverdines in the culture supernatants were determined by measuring integrated fluorescence emission between 445-460 nm after excitation at 420 nm [29,30]. The values were expressed relative to cell dry weight per ml.

### Survival and stress experiments

*P. extremaustralis* was cultured for 48h under microaerobic conditions in E2 cultures supplemented with *myo*-inositol or glucose and KNO_3_ (0.8 g/l). Cultures were exposed to 100 µM GSNO (m-NS) for 1h and cell viability was determined in plate assay.

Growth inhibition in response to oxidative or combined nitro-oxidative stress was evaluated by filter disk assay by growing *P. extremaustralis* in E2 medium supplemented with glucose or *myo*-inositol and increasing KNO_3_ concentrations of 0.8, 1.5 and 2.5 g/l that led to nitrite accumulation. Control cultures without nitrate were performed (oxidative stress only). LB plates were seeded with the different cultures and Whatman n° 1 filter discs (6 mm) were impregnated with 5 μl of 30% (v/v) H_2_O_2_ (Merck) as previously described [31]. Plates were incubated overnight, and the diameter of the halo was determined using the software ImageJ [32]. Nitrite concentration in the supernatant of cultures was determined following the method described by [33] and values were normalized to cell dry weight.

### Data availability

RNA-seq data were deposited in the European Molecular Biology Laboratory under accession number E-MTAB-11689.

### Statistical analysis

Differences between means were determined through the Student’s t test with confidence levels at > 95% in which P < 0.05 was considered as statistically significant. A Fisher’s exact test was performed to analyze the nitrosylation targets proteins. 1 or 2-ways ANOVA with multiple comparison were used when correspond.

## Results

### Effects of GSNO exposure in *P. extremaustralis*

Nitrosative stress effect on *P. extremaustralis* survival and growth under microaerobic conditions was evaluated using different S-nitrosoglutation (GSNO) concentrations. Growth in presence of different concentrations of GSNO was similar to the control culture except for 100 µM that showed a significant decrease (P<0.05) at the stationary phase (Fig. S2). Survival assays showed that when the bacterial culture was exposed to GSNO for 1h a significant decrease in bacterial viable counts (CFU/ml) was observed comparing with the control culture and provoked a 75% drop in survival in microaerobic cultures (Fig. 1). These results indicate that GSNO exposure affected *P. extremaustralis* survival in short time, thus we chose 100 µM GSNO (m-NS) exposure for 1h for further experiments.

**Figure 1.**
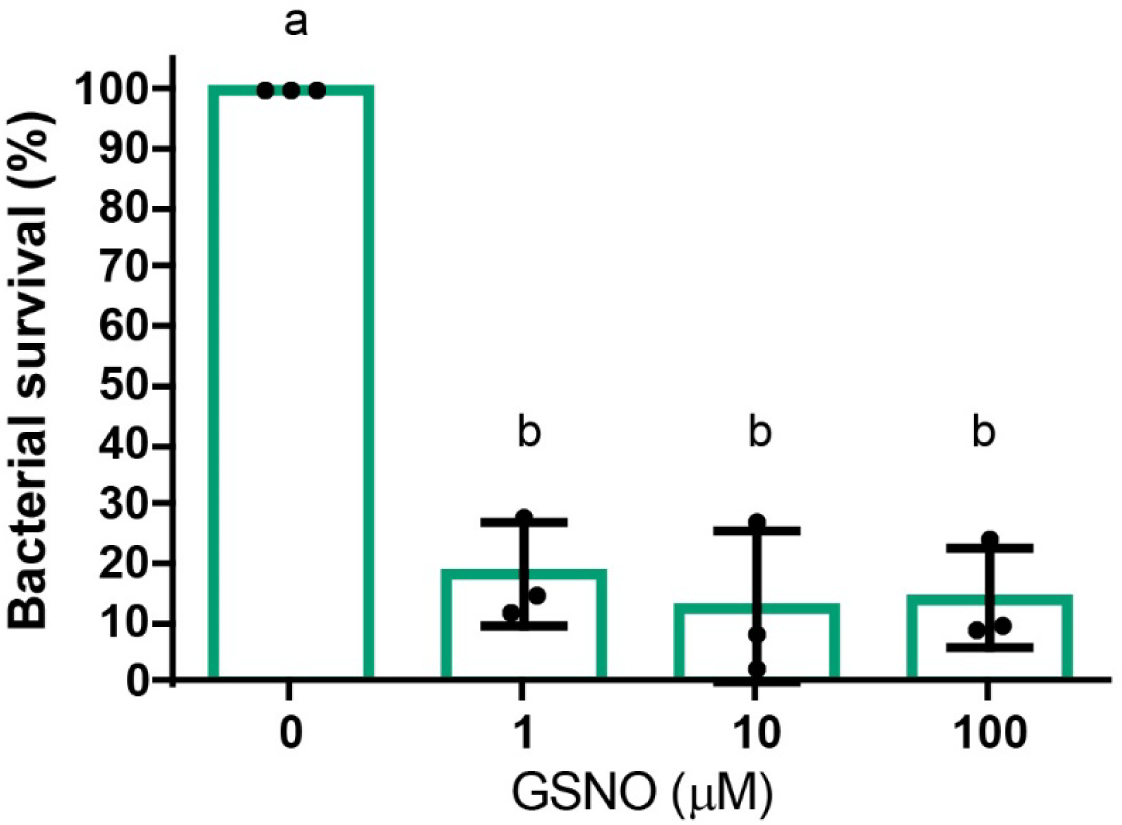
Survival of microaerobic cultures to GSNO exposure. Cultures were exposed during 1h and survival in LB plates was determine. Circles represent individual values of independent experiments. Error bars represent the standard deviation of the mean. Different letters denote significant differences using 1-way ANOVA with Tukey’s multiple comparisons test (P < 0.05)

### Transcriptomic profile of *P. extremaustralis* under low aeration and nitrosative stress conditions

*P. extremaustralis* response to nitrosative stress was evaluated by comparing transcriptomic data obtained by RNA sequencing (RNA-Seq) in microaerobic cultures m-C and m-NS. RNA-seq expression profile revealed 5888 transcripts. Rockhopper analysis showed 249 transcripts with differential expression in m-NS compared to m-C (P <0.05, Q <0.05). Among them, 86 genes were found to be repressed and 163 over-expressed, representing 1.46% and 2.77% of the total, respectively (Fig. S3a, Table S1). Several genes with differential expression did not have defined associated functions and were classified as “hypothetical”, representing 11.6% and 21% of the over-expressed and repressed genes, respectively. Among genes with a known predicted function, *P. extremaustralis* under m-NS showed an increased expression of genes involved in carbon, RNA, DNA and iron metabolism (Table S1).

RNA-seq results were validated by analyzing the expression of three selected genes, *motB*, *flaB*, and *hbo* using qRT PCR (Fig. S2b). In concordance with the RNA-seq results, we found higher expression levels of *motB,* encoding a flagellar motor rotation protein, in m-NS vs. m-C, whereas we observed a decrease in *hbO,* encoding the hemoglobin-like protein HbO (Fig. S2b, Table S1) and no differences in *flaB* expression between conditions.

### *P. extremaustralis’* response to nitrosative stress

To elucidate possible functional relationships, differentially expressed genes were categorized into functional groups (Fig. 2a). GSNO exposure caused increased expression of genes encoding different RNS detoxifying enzymes such as an iron-dependent superoxide dismutase (PE143B_0125285) and an alkyl hydroperoxide reductase (*ahpC*), a thioredoxin (PE143B_0111145) and a glutathione S-transferase (PE143B_0109905). A gene encoding another glutathione S-transferase (PE143B_0100180) was repressed in m-NS along with *hbO* (Fig. 2a, Table S1).

**Figure 2.**
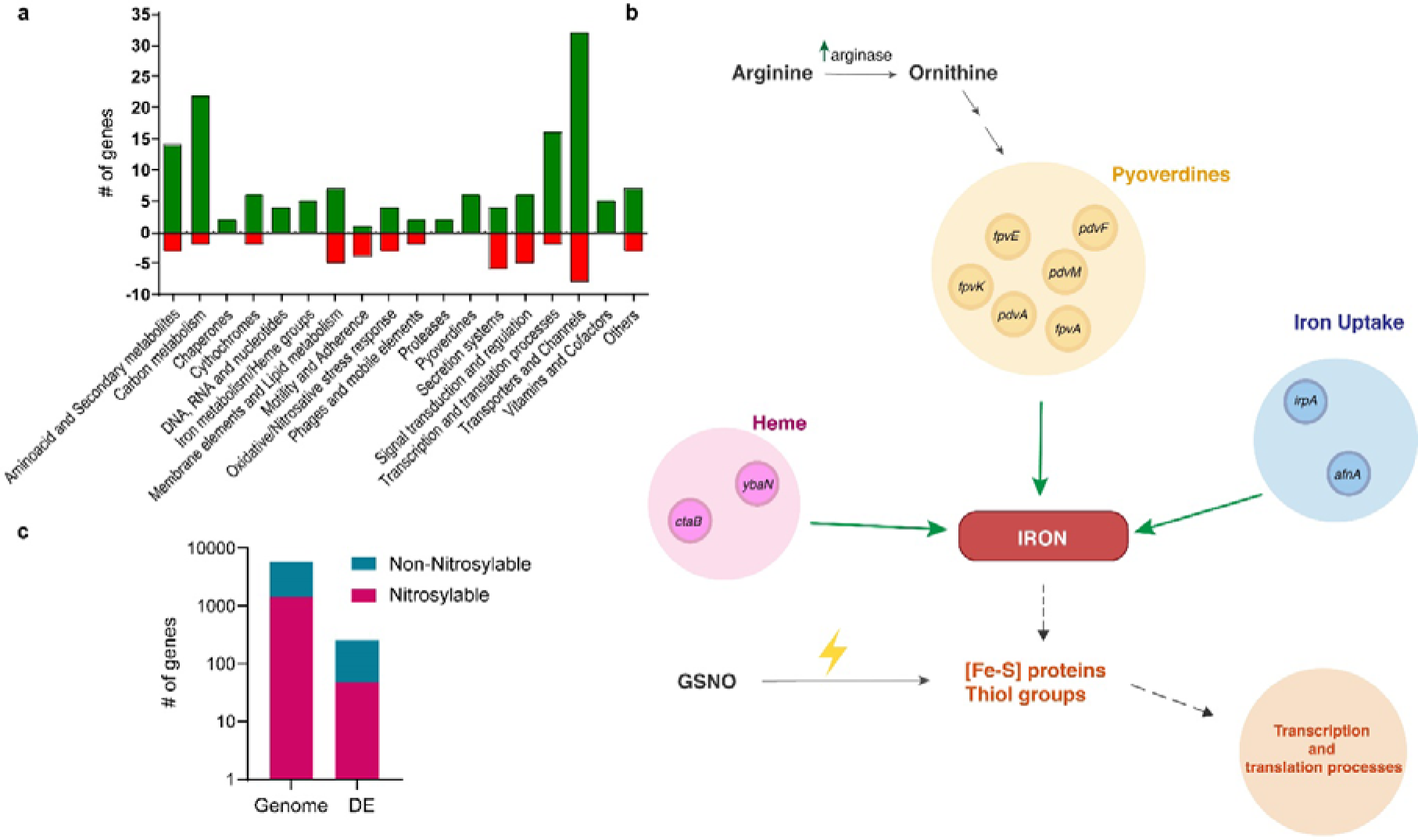
Effect of GSNO on gene expression profile. **a.** Classification of differentially expressed genes after GSNO exposure in microaerobiosis (m-NS vs. m-C) into functional categories. Hypothetical transcripts (with no inferred function) are not displayed. Green and red bars represent up- and down-regulated genes, respectively. **b.** Over-expressed genes related to iron metabolism. **c**. Nitrosylable proteins determined with Target Pathogens in the differential expressed genes compared to total nytrosilable proteins in *P. extremaustralis* genome. DE: differentially expressed genes. Fisher’s exact test, P = 0.0323.

Transcriptomic analysis showed an increase in mRNA expression of iron and heme transporters coding genes such as *ctaB, ybaN, irpA* and *afuA* (Fig 2b, Table S1). In m-NS we found upregulation in genes related to pyoverdine siderophore biosynthesis, a key function for iron uptake, including *pdvA, pdvM, pdvF, fpvA, fpvE* and *fpvK* (Fig. 2b, Table S1). Moreover, the arginase coding gene, was also upregulated in m-NS comparing with m-C. Arginase catalyzes the conversion of arginine to ornithine, which can then serve as a substrate for pyoverdine biosynthesis (Fig. 2b, Table S1).

Iron and other metals are key components of iron-sulfur proteins that could be target of RNS, leading to protein inactivation and triggering cellular signaling [34]. Therefore, we analyzed other cellular functions related with transcription and protein turn-over. GSNO treated cultures showed an increase in expression of RNA helicase (PE143B_0107010), RNA polymerase associated protein RapA, transcriptional terminator *nusB* (PE143B_0123570) and 11 ribosomal proteins coding genes (Table S1). In concordance, a gene encoding translation initiation inhibitor was repressed (Table S1). To deeply analyze the protein turnover related with RNS attack we performed an analysis using Target Pathogen platform to detect genes encoding proteins with free tyrosine or cysteine as well as metal binding sites which are reactive to RNS [1]. Among differentially expressed genes in m-NS we found that nitrosylable proteins were over-represented in comparison to non-nitrosylable when total genes of *P. extremaustralis* were considered (Fisher’s exact test, P = 0.0323). Overall, our results suggest that nitrosative stress under microaerobic conditions provokes an increase in the expression of genes involved in transcription and translation processes, probably to compensate the damage to proteins with Fe-S clusters or metal binding (Fig. 2c).

### Inositol metabolism in *P. extremaustralis*

Remarkably, RNAseq results showed increased expression of genes related to inositol catabolism which involves enzymes that convert inositol into acetyl-CoA and dihydroxyacetone phosphate (DHAP) (Fig. 3a, Table S1).

**Figure 3.**
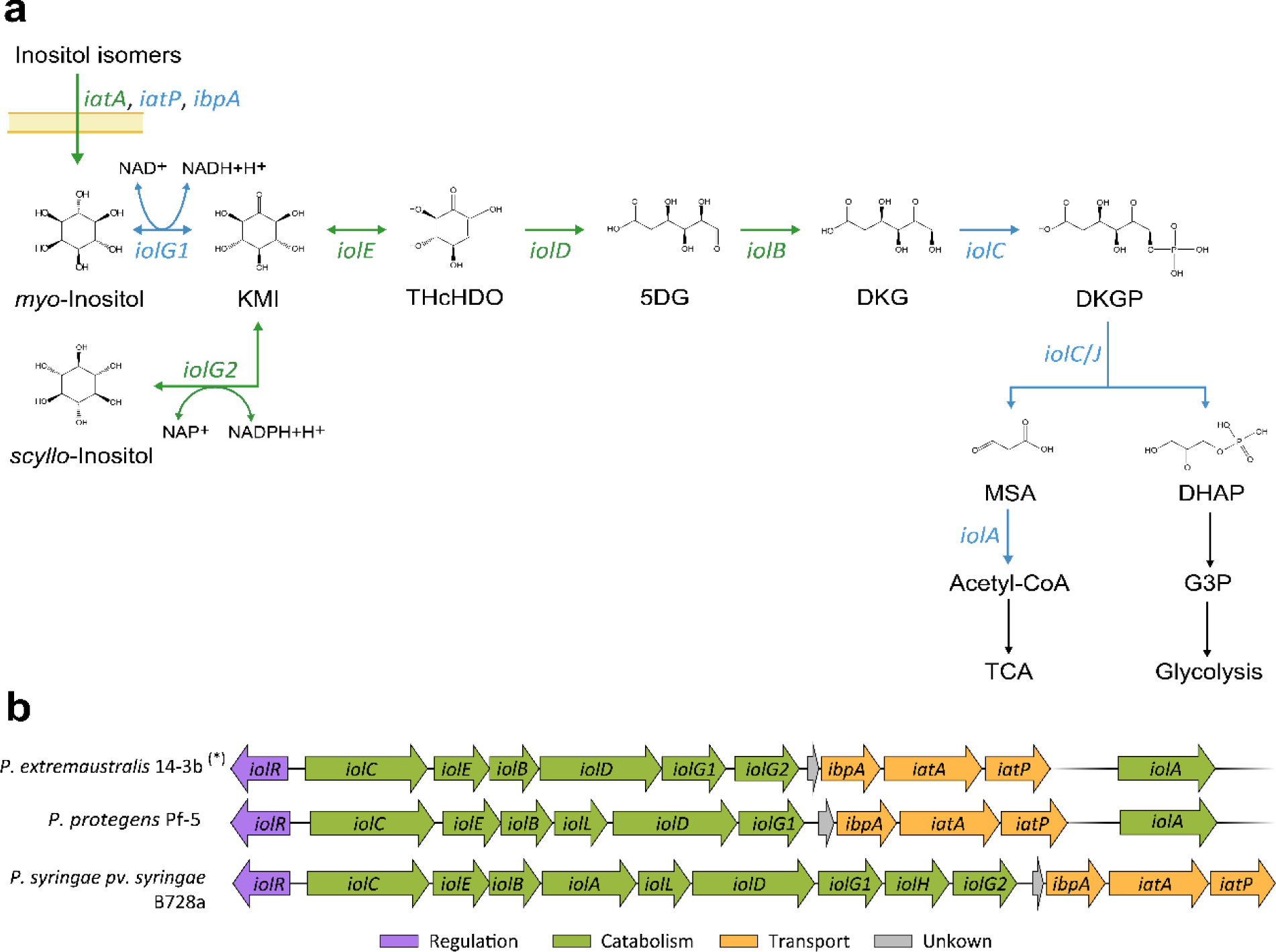
Metabolic pathway associated with inositol catabolism in *Pseudomonas extremaustralis* 14-3b: Expression under nitrosative stress and genetic organization. **a**. Gene expression and metabolic route related to inositol catabolism under m-NS Genes and arrows shown in green indicate up-regulated functions under m-NS, whereas those in blue represent functions with no expression differences between both conditions. **b**. Analysis of genes involved in inositol in *P. extremaustralis* in comparison with *P. protegens* Pf-5 and *P. syringae* pv. syringae B728a. (*) The same genetic organization was observed in USBA 515, DSM17835^T^, 2E-UNGS, CSW01,1906 and NQ5 *P. extremaustralis* strains. Compounds: KMI, 2-keto-*myo*-inositol; THcHDO, 3, 3D-(3,4/5) trihydroxycyclohexane-1,2-dione; 5DG, 5-deoxy glucuronic acid; DKG, 2-deoxy-5-keto-d-gluconic acid; DKGP, DKG 6-phosphate; DHAP, dihydroxyacetone phosphate; MSA, malonic semialdehyde; acetyl-CoA, acetyl coenzyme A; DHAP, dihydroxyacetone phosphate; G3P, glyceraldeyde-3-phosphate; TCA, tricarboxylic acid cycle. Genes: *iolG1*, *myo*-inositol dehydrogenase; *iolG2*, *scyllo*-inositol dehydrogenase; *iolE*, 2KMI dehydratase; *iolD*, THcHDO hydrolase; *iolB*, 5DG isomerase; *iolC/J*, DKG kinase + aldolase; *iolA*, MSA dehydrogenase; *iatA*, sugar ABC transporter ATP-binding protein; *iatP*, ABC transporter permease; *ibpA*, sugar ABC transporter substrate-binding protein.

Our results showed a significant up-regulation (P and Q<0.05) in mRNAs encoded in the *iol* operon in response to GSNO stress, including *iolB* (2-fold), *iolD* (1.75-fold), *iolE* (1.83-fold) *iolG2* (also known as *iolW* or *mocA*) (1.86-fold), *iolR* (2.62-fold), and the *iatA* gene (1.75-fold) (Fig. 3a; Table S1). No significant differences were observed in the expression of *iolA* (1-fold)*, iolG1*(1.54-fold) and *iolC/J* (1.25-fold) in both conditions, neither for *iatP* (1.20-fold) and *ibpA* (1.44-fold) (Fig. 3a). It was found that *iolC* possesses the DUF2090 domain that was recently proposed to encode 2-deoxy-5-keto-gluconic acid-6-phosphate aldolase [35]. This domain fulfills function originally attributed to *iolJ* which is not present in *P. extremaustralis* genome (Fig. 3b).

When we compared the inositol catabolic cluster of *P. extremaustralis* with *P. protegens* Pf-5, a well-known plant growth promoter species, and *P. syringae* pv. syringae B728a, a plant pathogen, we found some differences in this genomic region. Unlike *P. protegens* Pf-5, the genome of *P. extremaustralis* did not exhibit the presence of *iolL*. Conversely, we identified a duplication of *iolG* within this bacterial species. *P. syringae* pv. syringae B728a presented *iolH*, which is absent in both *P. protegens* and *P. extremaustralis*. Similar to *P. extremaustralis, P. syringae* pv. syringae B728a presented two copies of *iolG* but separated by *iolH* (Fig. 3b). The *iolA* gene in *P. extremaustralis* is located in a different genomic region like in *P. protegens* and other analyzed *Pseudomonas* [36]. However, in P. *syringae* pv. syringae B728a is included in the *iol* catabolic cluster (Fig. 3b). Furthermore, we found that the *iol* gene cluster of *P. extremaustralis* exhibited a high degree of intraspecific conservation showing the same organization as 14-3b in all genomes analyzed, including strains USBA 515, DSM17835^T^, 2E-UNGS, CSW01,1906, NQ5.

As we detected the presence and expression of *iol* genes, we first tested *P. extremaustralis* capability to grow under aerobic conditions using *myo*-inositol as sole carbon source. Growth was similar between *myo*-inositol and glucose supplemented cultures reaching an OD_600nm_ value of 2.02 ± 0.15 and 2.22 ± 0.15, respectively. Under microaerobic conditions *P. extremaustralis* developed an evident biofilm in *myo*-inositol supplemented cultures (Fig. 4a). Therefore, OD_600nm_ and cellular dry weight (DW) were determined (Fig. 4b). Similar results were obtained for both carbon sources regardless of the approach employed for growth estimation. Biofilm formation was assayed in E2 medium supplemented with glucose or inositol. In concordance, we also found that with *myo*-inositol supplemented cultures *P. extremaustralis* showed a higher biofilm index compared to glucose (Fig. 4c).

**Figure 4.**
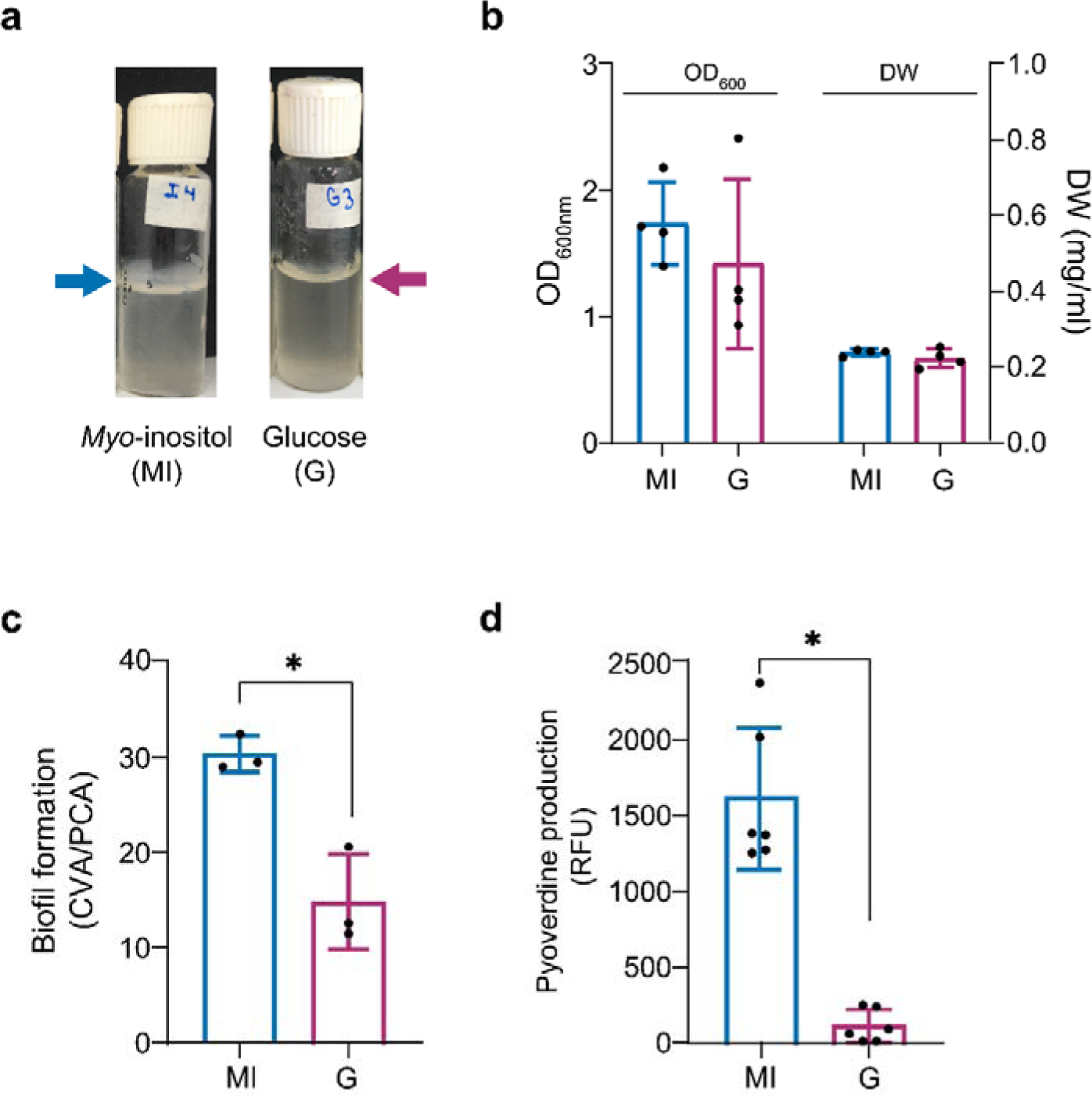
Physiological features of *P. extremaustralis* using *myo*-inositol o glucose as carbon source. **a.** Photograph of microaerobic *myo*-inositol or glucose supplemented cultures showing the biofilm in the surface. **b.** Microaerobic growth measured by optical density (OD_600nm_) or Cellular dry weight (DW). **c.** Biofilm formation index (ACV/APC) in *myo*-inositol or glucose. **d.** Effect of carbon sources on pyoverdine production. RFU: Relative Fluorescence Units. Fluorescence units were normalized to the cell dry weight/ml. Error bars represent the standard deviation of the mean.

Additionally, considering the upregulation of coding genes related with pyoverdine biosynthesis after treatment with GSNO, we investigated siderophores production in *P. extremaustralis* using *myo*-inositol as carbon source. Pyoverdine production was higher for *myo*-inositol supplemented cultures comparing to glucose (Fig. 4d)

### *Myo*-inositol impact on stress response

Further, we investigated stress resistance when *myo*-inositol was used as sole carbon source under low oxygen conditions. To analyze the nitrosative stress response, we performed a survival test using microaerobic cultures in E2 medium supplemented with *myo*-inositol or glucose exposed to GSNO treatment. We found no differences in survival between those treated with GSNO and control cultures despite of the carbon source (Fig. 5a).

**Figure 5.**
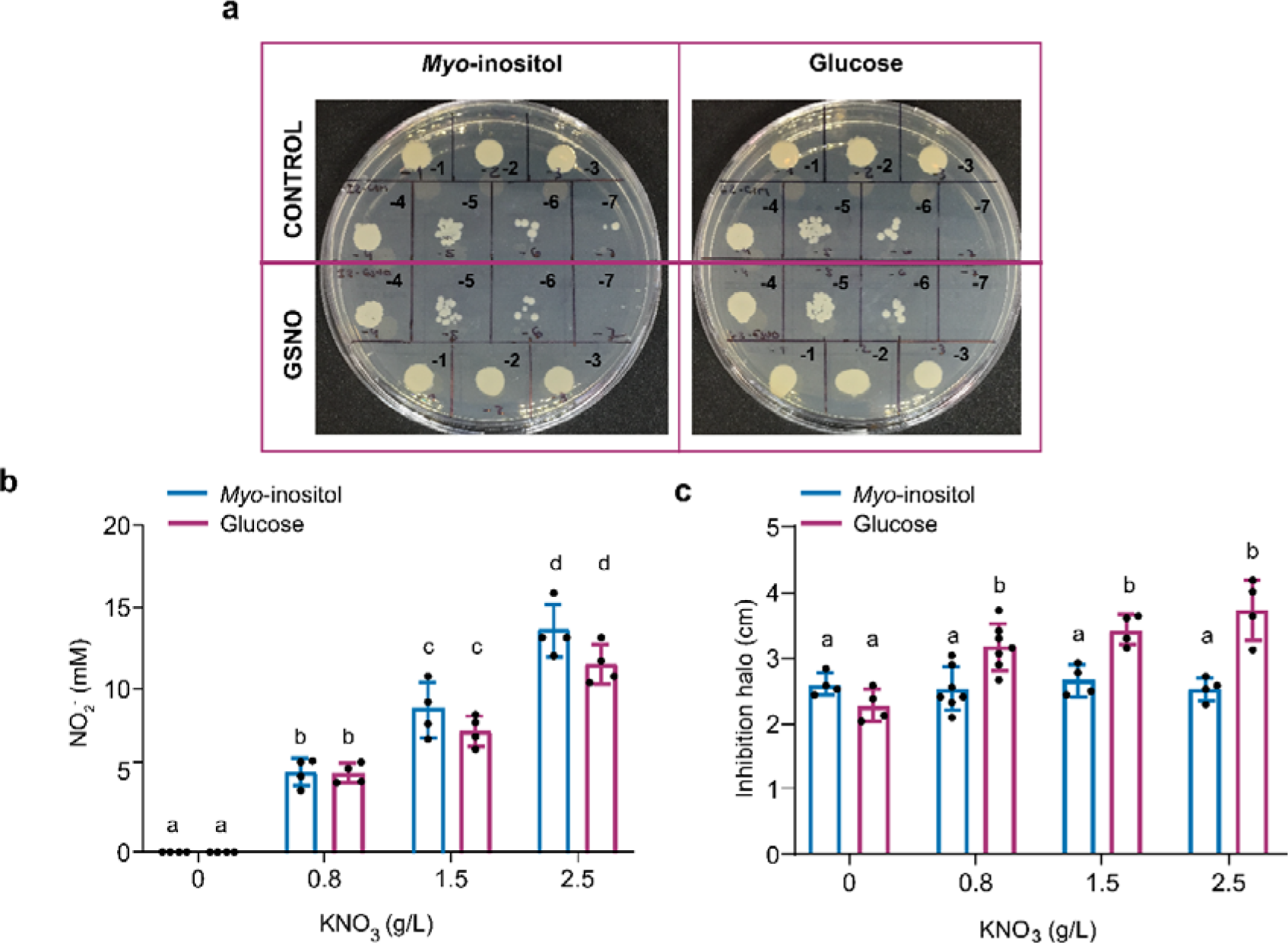
Nitrosative and combined nitro-oxidative stress resistance in *P. extremaustralis*. **a.** Survival after GSNO exposure. **b.** Nitrite production in cultures supplemented with *myo*-inositol or glucose and different nitrate concentrations. **c.** Inhibition halos of cultures performed in (**b**) exposed to H_2_O_2_ (Combined nitro-oxidative stress). Error bars represent the standard deviation of the mean.

*P. extremaustralis* is only capable to reduce nitrate to nitrite which accumulates in the extracellular medium. Therefore, the resistance to nitro-oxidative stress derivate from the combination of the nitrite accumulated and H_2_O_2_ was analyzed in *myo*-inositol and glucose supplemented cultures, following the scheme shown in Fig. S4. Nitrite production was detected in cultures supplemented with *myo*-inositol and glucose as sole carbon source and different nitrate concentrations reaching a maximum of 13.553± 1.624, confirming also the *myo*-inositol utilization in a respiratory pathway (Fig. 5b). When *P. extremaustralis* was grown in *myo*-inositol or glucose supplemented media without nitrate, the oxidative stress resistance was similar for both carbon sources while the nitro-oxidative stress resistance was higher for cultures grown in *myo*-inositol in all tested KNO_3_ concentrations (Fig. 5c).

## Discussion

Bacterial species are subjected to different stress conditions including lack of nutrients, changes in temperature or oxygen availability and oxidative and nitrosative stress, among others. Oxidative and nitrosative stress can be produced by the imbalance between the ROS or RNS generation and the incapability to detoxify them [37].

In response to these stress conditions, bacteria may activate various response mechanisms, such as the production of chaperons and detoxicant enzymes, the adjustment of gene expression, and the modification of their cell envelope composition. A common effect during stress is the reprograming of transcription and translation processes including the reduction of ribosome biogenesis, the modification of the translational machinery and the regulation of initiation and elongation of certain proteins [38]. Interestingly, gene expression during stress involves not only the transcription processes but also the modification of mRNA stability and in overall with translation provokes a transcriptome stability [39,40]. In this work, we found an up-regulation of the transcription and translation machinery probably to compensate the damage caused by RNS particularly on nitrosylable proteins. The Fe-S clusters, that can be found in three different forms, exhibit high capacity of accepting or donating electrons thus are important for redox response and act as redox sensors like the Fnr regulator in *Escherichia coli* or Anr in *Pseudomonas* species [41–43].These proteins and those containing metals are involved in several essential cellular functions like respiration, central carbon catabolism and RNA and DNA processes and are a target of RNS damage [41] Particularly, it has been reported that the reaction between nitric oxide (NO) with Fe-S clusters lead to protein degradation and breakdown of the cluster generating dinitrosyl iron complexes [44]. Our analysis using Target Pathogen platform showed the enrichment of genes encoding nitrosylable proteins in the differentially expressed ones, particularly in those upregulated. Our hypothesis is that to cope with GSNO derived stress the Fe-S and metal binding proteins need to be replaced. This also could explain the upregulation of Fe-uptake related genes observed in presence of GSNO. Iron is involved in the Fenton reaction in presence of H_2_O_2_, derived from endogenous or exogenous sources, that could exacerbate the oxidative damage but paradoxically this metal is also necessary for Fe-S and metal protein biogenesis [45,46]. This complex scenario additionally requires the involvement of enzymes and antioxidant cycles. In presence of GSNO we found upregulated the coding genes of glutathione S-transferase, superoxide dismutase (SOD), thiol-disulfide isomerase, thioredoxins and alkyl hydroperoxide reductase subunit C-like protein. The alkyl hydroperoxide reductase, important during non-lethal stress, SOD as well as the oxidation of thioredoxins in response to oxidative stress depends on NAD(P)H availability and in overall reduction power is needed for the essential process for survival and to cope with stress [47]. Interestingly, the needed of NADH and/or NADPH production during stress could represent a challenge for bacterial cells. Glycolysis and TCA provides the NADH and energy necessaries for bacterial survival but also creates an oxidative scenario through the respiration chain [48]. It has been reported several metabolic strategies to obtain energy and reducing power besides glycolysis such as aminoacid catabolism, ethanol oxidation and in *P. fluorescens* an entire metabolic reprogramming has been reported in presence of oxidative stress to generate NADPH [49]. In this previous work the authors showed an increase in the activity and expression of malic enzyme (similar to this work in presence of GSNO), glucose-6-phosphate dehydrogenase and NADP+-isocitrate dehydrogenase [49].

In addition, we found that in presence of GSNO under microaerobic conditions *P. extremaustralis* over-expressed several *iol* catabolic genes including the transcriptional regulator and the *iatA* gene. We also demonstrated that this bacterium was effectively capable of using inositol as its sole carbon source. Inositol is a sugar alcohol present in all domains of life and presents various structural isomers [50], with *myo*-inositol being the most abundant in nature. *Myo*-inositol is found in high quantities in plants, including root exudates, and may also originate from the dephosphorylation of phytate, one of the major phosphorus storage molecules for plants [51]. Inositol catabolism has been experimentally demonstrated in several bacterial species, both Gram positive and Gram negative, belonging to genera such as Rhizobium*, Lactobacillus, Klebsiella, Corynebacterium, Legionella, Bacillus, and Salmonella* [52–58]. Furthermore, an extensive analysis of inositol catabolic genes in thousands of bacterial genomes revealed a widespread distribution within different ecological niches [51].

While inositol metabolism in bacteria has garnered recent attention, its role in stress resistance remains unexplored. Remarkably, our findings revealed that inositol protects against nitro-oxidative stress, and increases biofilm and pyoverdine production, which are relevant traits for environmental adaptability. Inositol catabolism leads to the production of glyceraldehyde-3 phosphate, and acetyl-CoA both intermediaries of central energy generation metabolism. In addition, *iolG2,* which was found overexpressed after GSNO exposure, is related to NAPH generation, necessary for the function of antioxidative defenses.

It is worth noting that previous transcriptome analysis of *P. extremaustralis* performed under cold conditions or under microaerobic conditions with or without oxidative stress did not show the upregulation of inositol catabolic genes [14,59,60], indicating a fine-tuning response to nitrosative stress.

Some RNS also play a crucial role in cell signaling. For instance, in plants, NO mediates for various processes including root hair growth, stomatal closure, programmed cell death and other responses that help them adapt to changing environmental conditions [8] whereas in bacteria, NO plays a role as intermediary in denitrification processes, but also in biofilm formation and quorum sensing [61]. The *iol* gene cluster was identified in both plant-pathogenic bacteria, and rhizospheric plant growth promoting bacteria forming symbiotic root nodules [51].

Recently, the *iol* gene cluster in *Pseudomonas* was identified as an important trait in root colonizers, by increasing swimming motility and siderophore production in response to plant derived inositol [36]. Increased pyoverdine siderophore production in presence of *myo*-inositol was also observed in this work. *P. extremaustralis* possesses plant growth-promotion traits, including the ability to solubilize phosphate and produce indole acetic acid [62]. Since NO is also a signaling molecule released by plants [63], we hypothesize that NO released by GSNO mimic the NO production of plants, triggering the differential expression of the inositol metabolic pathway conferring an adaptive advantage.

In conclusion, our findings reveal a multifaceted and finely tuned response mechanism to nitrosative stress that encompasses transcriptional reprogramming, the upregulation of genes encoding key proteins involved in iron homeostasis, and the activation of inositol catabolism. Notably, experimental results point to inositol metabolism as a novel mechanism to cope with stress.

## Acknowledgments

This work was funded by grants from Universidad de Buenos Aires and Consejo Nacional de Investigaciones Científicas y Técnicas (CONICET). We are thankful to Dr. Jörg Vogel (Institut für Molekulare Infektionsbiologie (IMIB) – University of Würzburg) for his guidance and for providing resources for conducting the RNAseq experiments. E.C.S.V. received a Deutscher Akademischer Austauschdienst (DAAD) fellowship for short-term research stays at the RNA biology group (IMIB – University of Würzburg, Germany).

