## Supplementary Figures for "Nitrosative stress under microaerobic conditions triggers inositol metabolism in *Pseudomonas extremaustralis*"

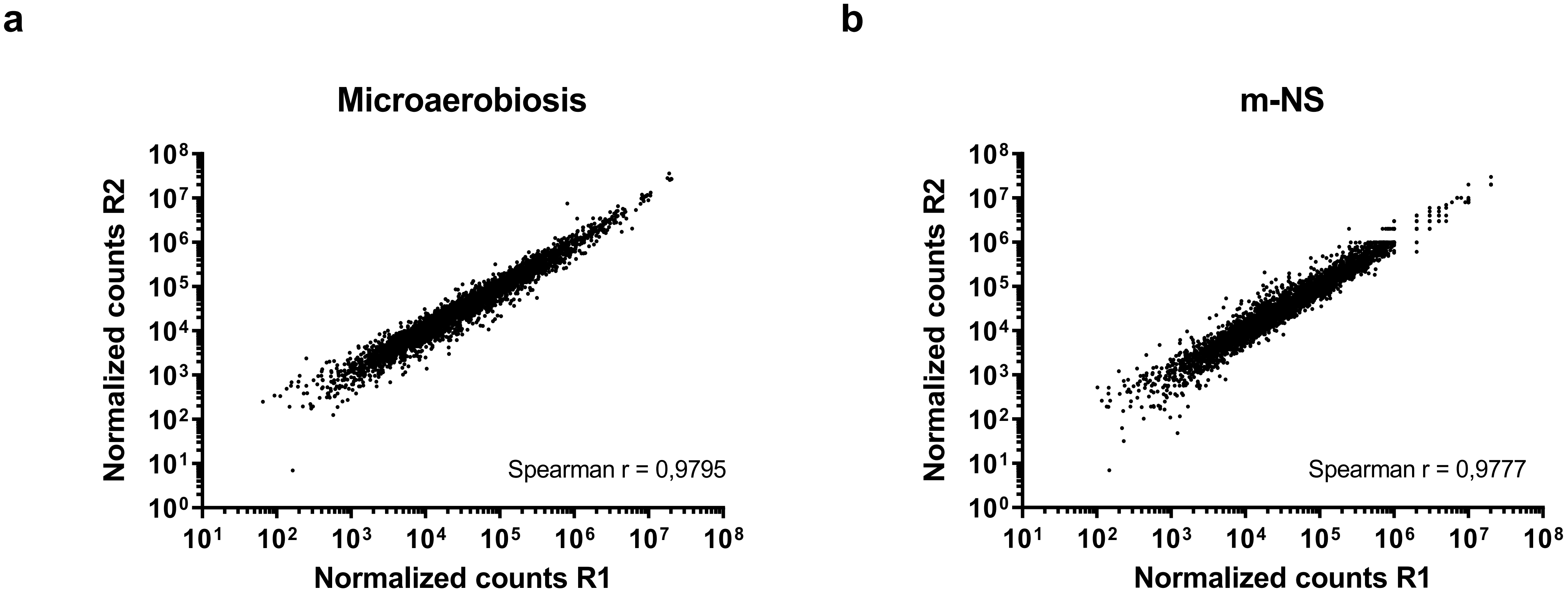


Figure S1: Dot plot representing normalized counts for each RNA-seq replicate. a. Microaerobic culture conditions. b. Microaerobic culture subjected to GSNO (m-NS).

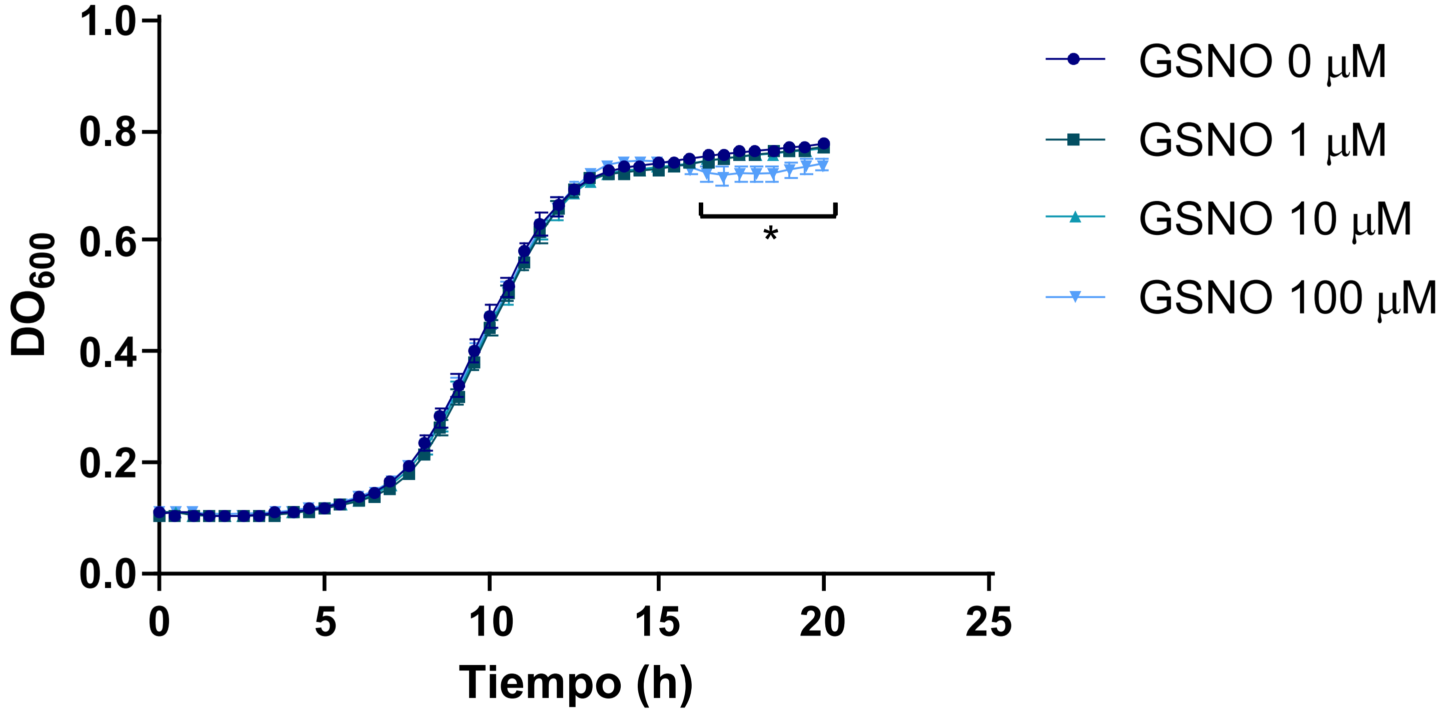


Figure S2: Growth curves of *P. extremaustralis* in microaerobiosis with different GSNO concentrations. Values represent the mean ± SD of 6 independent experiments.

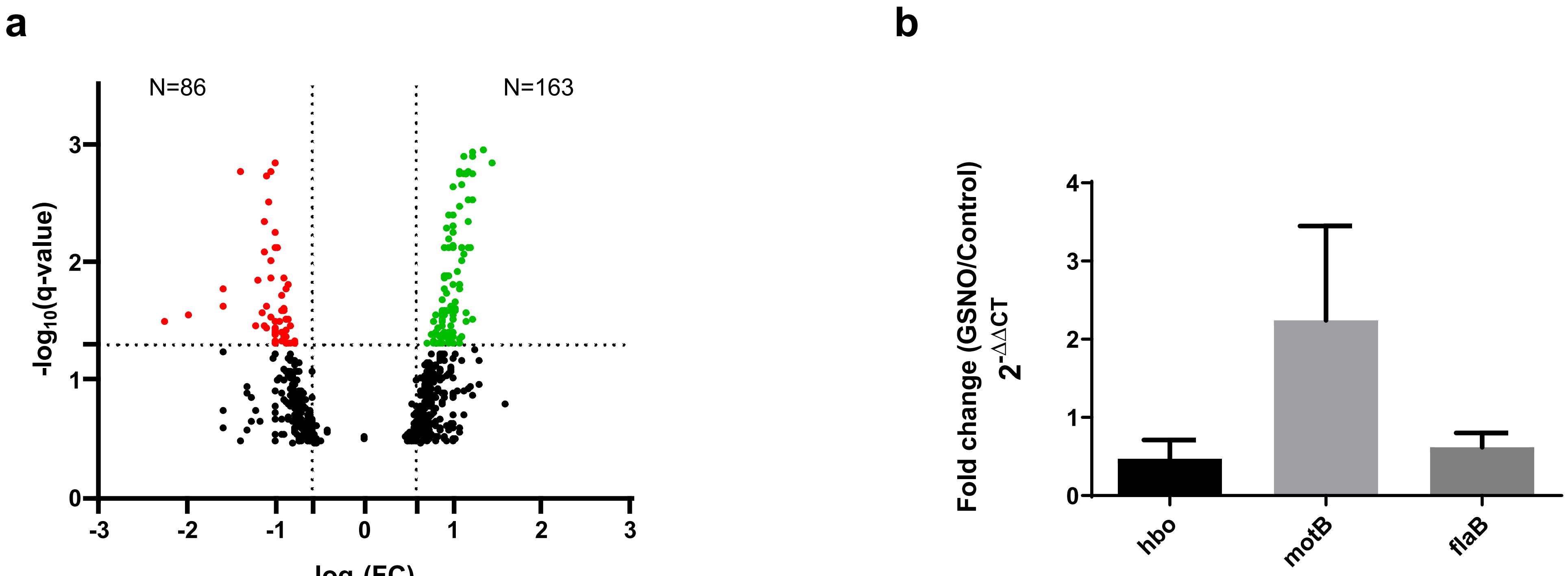


Figure S3 a. Volcano plots depicting –log10 (Q-value) versus log2 (Fold Change -FC-).Differentially expressed genes (DEGs) are represented by colored dots. Green dots represent upregulated genes (P-value and Q-value <0.05, Fold change >1.5) and red dots represent downregulated genes (P-value and Q-value <0.05, Fold change < -1.5). For the construction of the Volcano plots, genes were initially filtered based on their P-values. All resulting genes were plotted and filtered again by Q-value. b. Quantitative Real Time PCR. Comparative expression analysis of 3 selected genes between m-NS and microaerobic growth conditions. Values represent the mean ± SD of three independent experiments.

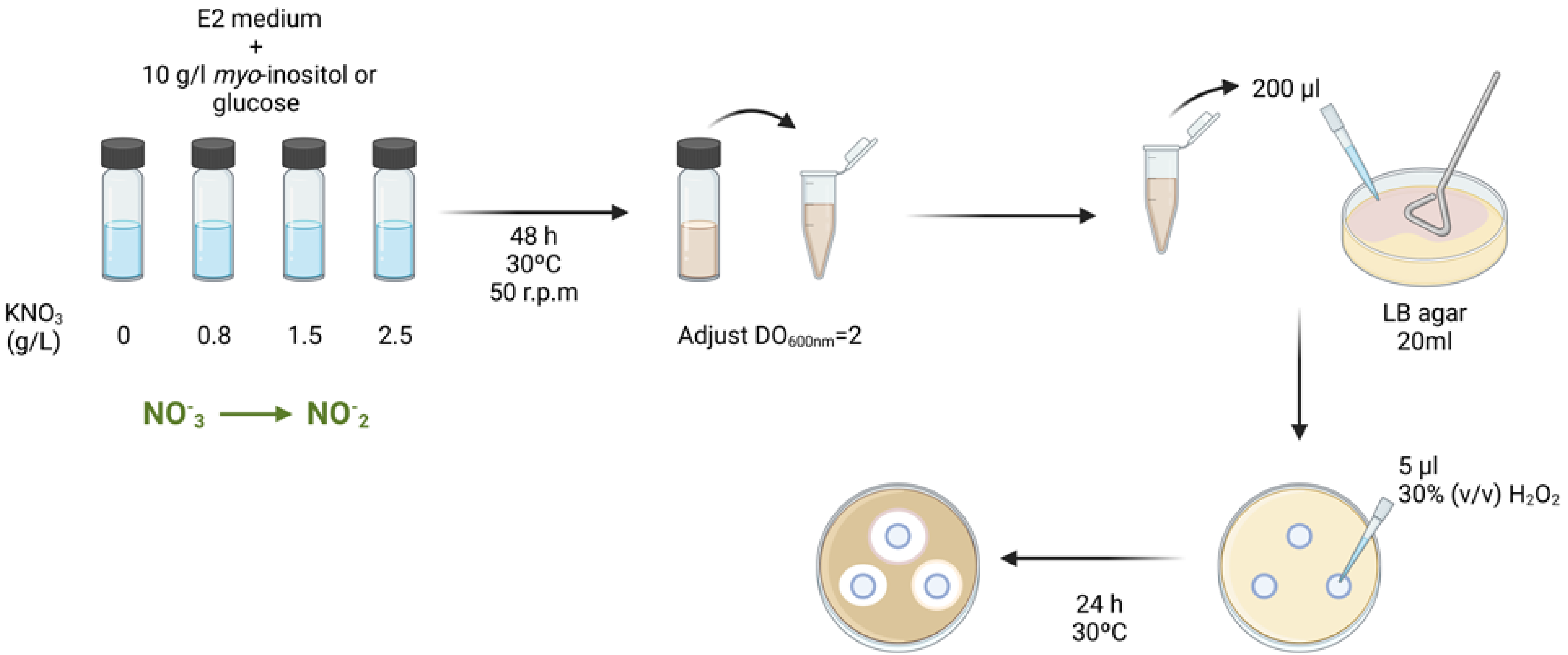


Figure S4: Scheme of the assay to study the resistance to nitro-oxidative stress derivate from the combination of the nitrite accumulated and $H_2O_2$.
